# SARS-CoV-2 genome sequencing with Oxford Nanopore Technology and Rapid PCR Barcoding in Bolivia

**DOI:** 10.1101/2021.07.06.451357

**Authors:** Oscar M. Rollano-Peñaloza, Carmen Delgado Barrera, Aneth Vasquez Michel

## Abstract

SARS-CoV-2 genomic surveillance has Illumina technology as the golden standard. However, Oxford Nanopore Technology (ONT) provides significant improvements in accessibility, turnaround time and portability. Characteristics that gives developing countries the opportunity to perform genome surveillance. The most used protocol to sequence SARS-CoV-2 with ONT is an amplicon-sequencing protocol provided by the ARTIC Network which requires DNA ligation. Ligation reagents can be difficult to obtain in countries like Bolivia. Thus, here we provide an alternative for library preparation using the rapid PCR barcoding kit (ONT). We mapped more than 3.9 million sequence reads that allowed us to sequence twelve SARS-CoV-2 genomes from three different Bolivian cities. The average sequencing depth was 324X and the average genome length was 29527 bp. Thus, we could cover in average a 98,7% of the reference genome. The twelve genomes were successfully assigned to four different nextstrain clades (20A, 20B, 20E and 20G) and we could observe two main lineages of SARS-CoV-2 circulating in Bolivia. Therefore, this alternative library preparation for SARS-CoV-2 genome sequencing is effective to identify SARS-CoV-2 variants with high accuracy and without the need of DNA ligation. Hence, providing another tool to perform SARS-CoV-2 genome surveillance in developing countries.

## Introduction

Coronavirus (CoV) are virus with the largest known single-strand RNA genomes (Yin and Wunderink 2018). Coronavirus are enveloped virus with several trimeric-spikes on their surface which altogether form a crown or “corona” (V’kovski et al. 2021; Zhang and Kutateladze 2020).

Coronavirus can cause respiratory and enteric infections in humans and are one of the main agents of the severe acute respiratory syndrome (SARS) (Cheng et al. 2007). The first coronavirus that caused a mayor pandemic of SARS (SARS-CoV) emerged on November 2002 in Guangdong, China (Peiris et al. 2003; Skowronski et al. 2004). Ten years later, another mayor pandemic was caused by the Middle East respiratory syndrome-CoV (MERS-CoV) with its first patient reported on June 2012 in Jeddah, Saudi Arabia (Zaki et al. 2012). SARS-CoV and MERS-CoV infected more than ten thousand people with approximately 1600 deaths (Yin and Wunderink 2018). Both outbreaks marked the possibility of a SARS-CoV reemergence if we wouldn’t maintain the barriers between natural Coronavirus reservoirs and human society (Cheng et al. 2007; Cui et al. 2019).

All the conditions were finally met again on December 2019, when another Coronavirus (SARS-CoV-2) emerged on Wuhan, China (Zhou et al. 2020), causing the third and largest coronavirus outbreak. SARS-CoV-2 is the causal agent of the Coronavirus disease-19 (COVID-19) that has spread throughout the whole planet. SARS-CoV-2 has caused more than 3 million deaths worldwide at the time of writing (Ortiz-Prado et al. 2020). Coronavirus have higher mutation rates than DNA viruses which can result in a high viral genetic diversity and adaptability (Duffy 2018). The unstopped spreading of the virus allows opportunities for viral replication and mutations that have already driven the emergence of SARS-CoV-2 variants of concern, which may scape the scope of protection from vaccines and natural immunity.

Emergence of variants can drive the outcome of the pandemic in unpredicted ways. For example, in Manaus (Brazil) where more than 76% of the population was already infected and reached herd immunity (Buss et al. 2021), a new outbreak occurred by the appearance of a new SARS-CoV-2 lineage P1, also known as 501Y.V3 lineage inside the 20J clade (Sabino et al. 2021). Therefore, there is a need to surveil the emergence of new SARS-CoV-2 variants as frequent as possible and with all the tools available.

Although Illumina technology is the gold standard for SARS-CoV-2 genome sequencing Oxford Nanopore technology (ONT) has contribute to outbreak surveillance all over the world (Chiara et al. 2020), especially in countries with low GDP. For example, Rwanda a country with similar population to Bolivia and a GDP 4-times lower performs SARS-CoV-2 genome surveillance biweekly across the country (Butera et al. 2021).

The most used protocol to sequence SARS-CoV-2 with nanopore technology has been the SARS-CoV-2-v3-LoCost amplicon-sequencing protocol provided by the ARTIC Network (González-Recio et al. 2021; Tyson et al. 2020). Library preparation in this protocol requires DNA ligation, which can be cumbersome, and its reagents are difficult to get in countries like Bolivia. Thus, here we provide an alternative for library preparation using the rapid PCR barcoding kit (ONT) and we report the first SARS-CoV-2 genomes sequenced in Bolivia.

## Methods

### Sample collection

Purified RNA from nasopharyngeal samples positive for SARS-CoV-2 were remitted to Hospital San Pedro Claver, Chuquisaca, Bolivia. Sample information was removed to avoid patient identification except for sample location and technical data (Detection Cycle threshold (Ct) and extraction method). Sample selection criteria was high viral load (Ct<25).

RNA samples were obtained from three PCR diagnostic labs approved by the Health Minister of Bolivia to detect SARS-CoV-2. Hospital San Pedro Claver (HSPCl), Sucre; Laboratorio de referencia Departamental para la Vigilancia Epidemiológica – SEDES-Potosí (LDVEP), Potosí and Clin & Gen Lab, La Paz. RNA at HSPCl was extracted with the Viral Nucleic Acid Extraction Kit (IBI Scientific, IA, USA). RNA at Clin & Gen and LDVEP was extracted with the Viral RNA Extraction from Respiratory Specimens kit (Biomiga, CA, USA).

### Library preparation and Nanopore sequencing

RNA was reverse-transcribed using the RevertAid H minus First Strand cDNA Synthesis kit (Invitrogen, CA, USA) with random hexamer primers following the manufacturer’s instructions.

SARS-CoV-2 genome amplification was done based on Nanopore protocols described before for Ebola (Quick et al. 2016), Zika (Quick et al. 2017) and SARS-CoV-2 (Tyson et al. 2020). SARS-CoV-2 enrichment was done with the V3 primer pool (IDT, NJ, USA) designed by the ARTIC Network and Itokawa et al. (2020) using a Phusion High-Fidelity PCR Polymerase (Thermo Scientific, CA, USA). Multiplex PCR conditions were 1 cycle of: 98°C, 30 s; 30 cycles of 98° C, 15 s; 65°C, 5 min; 1 final cycle of 72°C for 7 min. For the protocol standardization we tried the following change in annealing and extension: 63°C, 1 min; 72°C, 4min.

Sample barcoding for the sequencing libraries was done with the Rapid PCR Barcoding Kit (SQK-RPB004) (Oxford Nanopore Technologies, United Kingdom) according to the manufacturer’s instructions with the following modifications: Fragmentation was done by 5 min at 30°C.

Sequencing libraries were quantified with Qubit (Thermo Scientific, CA, USA) and purified with the Magnetic beads AxyPrep: MAG PCR Clean-Up Kit (Axygen-Corning, AZ, USA). All barcoded samples were pooled together (~15ng DNA) and prepared to be loaded onto a R9.4.1 flowcell (Nanopore technologies, Oxford, UK) according to the manufacturer’s instruction. Samples were sequenced in a MinIon device for a total of 18 hours.

Basecalling was performed with Guppy v3.2.1 using the fast module. Consensus genomes were generated with the ARTIC Network bioinformatic pipeline (https://artic.network/ncov-2019/ncov2019-bioinformatic.sop.html) using the Medaka tool for calling variants.

Sequences whose depth was less than 10x were annotated as missing data (“Ns”) in the consensus sequences (Table 1). Consensus genome sequences were deposited at the GISAID database with the following accessions: EPI_ISL_1250842, EPI_ISL_1278275-1278284 and EPI_ISL_1363787.

**Table 1.** Nanopore sequencing genomic summary of 12 SARS-CoV-2 samples from Bolivia

| *Sample* | *Mapped Reads<sup>a</sup>* | *Depth* | *Length (bp)* | *Ns (Missing)* | *PANGO Lineage* |
| --- | --- | --- | --- | --- | --- |
| La Paz-1 | 233.733 | 389 X | 29.782 | 132 | B.1.1.348 |
| La Paz-2 | 332.735 | 390 X | 29.782 | 415 | B.1.1.348 |
| La Paz-3 | 141.075 | 272 X | 29.494 | 1.336 | B.1.1.348 |
| La Paz-4 | 243.159 | 363 X | 29.782 | 398 | B.1.177 |
| Sucre-1 | 634.401 | 320 X | 29.323 | 1.972 | B.1.1.348 |
| Sucre-2 | 447.325 | 331 X | 29.457 | 2.197 | B.1.1.274 |
| Sucre-3 | 432.514 | 286 X | 29.456 | 3.216 | B.1.2 |
| Sucre-4 | 465.285 | 296 X | 29.323 | 3.226 | B.1.1 |
| Potosi-1 | 398.489 | 339 X | 29.494 | 1.075 | B.1.1.348 |
| Potosi-2 | 208.357 | 301 X | 29.442 | 1.075 | B.1.1.274 |
| Potosi-3 | 145.561 | 327 X | 29.494 | 651 | B.1.1.274 |
| Potosi-4 | 276.495 | 340 X | 29.494 | 550 | B.1.1 |
| *Average* | 329.927 | 324,4 X | 29.526,9 | 1.353,58 |  |
| *Total* | 3.959.129 |  |  |  |  |

<sup>a</sup>Total reads mapped to the SARS-CoV-2 Wuhan-hu reference genome

### Genome assessment, variant identification and phylogenomic analysis

Genome consensus sequences from Bolivia were assessed and assigned to their respective PANGO lineages with Pangolin v3.1 version 2021-06-15. Nextstrain clades were assigned with the web app NextClade v1.4.0 (Hadfield et al. 2018). For the phylogenomic analysis, all complete SARS-CoV-2 genome sequences from neighboring regions of countries that surround Bolivia available on January, 2021 were downloaded from the GISAID repository (Acre, Mato Grosso, Mato Grosso do Sul and Rondônia from Brasil, and all sequences available from Argentina, Chile, Paraguay and Peru).

Genomic alignments were made using Muscle (Edgar 2004). Nucleotide and aminoacid substitution models were evaluated by MEGA X (Kumar et al. 2018). The phylogenetic tree was performed by maximum likelihood using the General Time Reversible model (Tavaré 1986). Total sequences in the exploratory dataset were 427 and the final dataset had 47 nucleotide sequences and had a total of 29903 positions (Table S1). Bootstrap testing was conducted with 50 replicates for exploratory analysis and with 1000 replicates for the final dataset. Bioinformatic analyses were computed at Centro Nacional de Computación Avanzada en Genómica y Bioinformatica (www.cncabo.umsa.bo).

## Results

This work is the first Coronavirus (SARS-CoV-2) genome sequencing carried out in Bolivia. We selected twelve samples of positive patients for Covid-19 from three different cities: 4 samples from Sucre, 4 samples from Potosí and 4 samples from La Paz (Fig. 1).

**Fig. 1.**
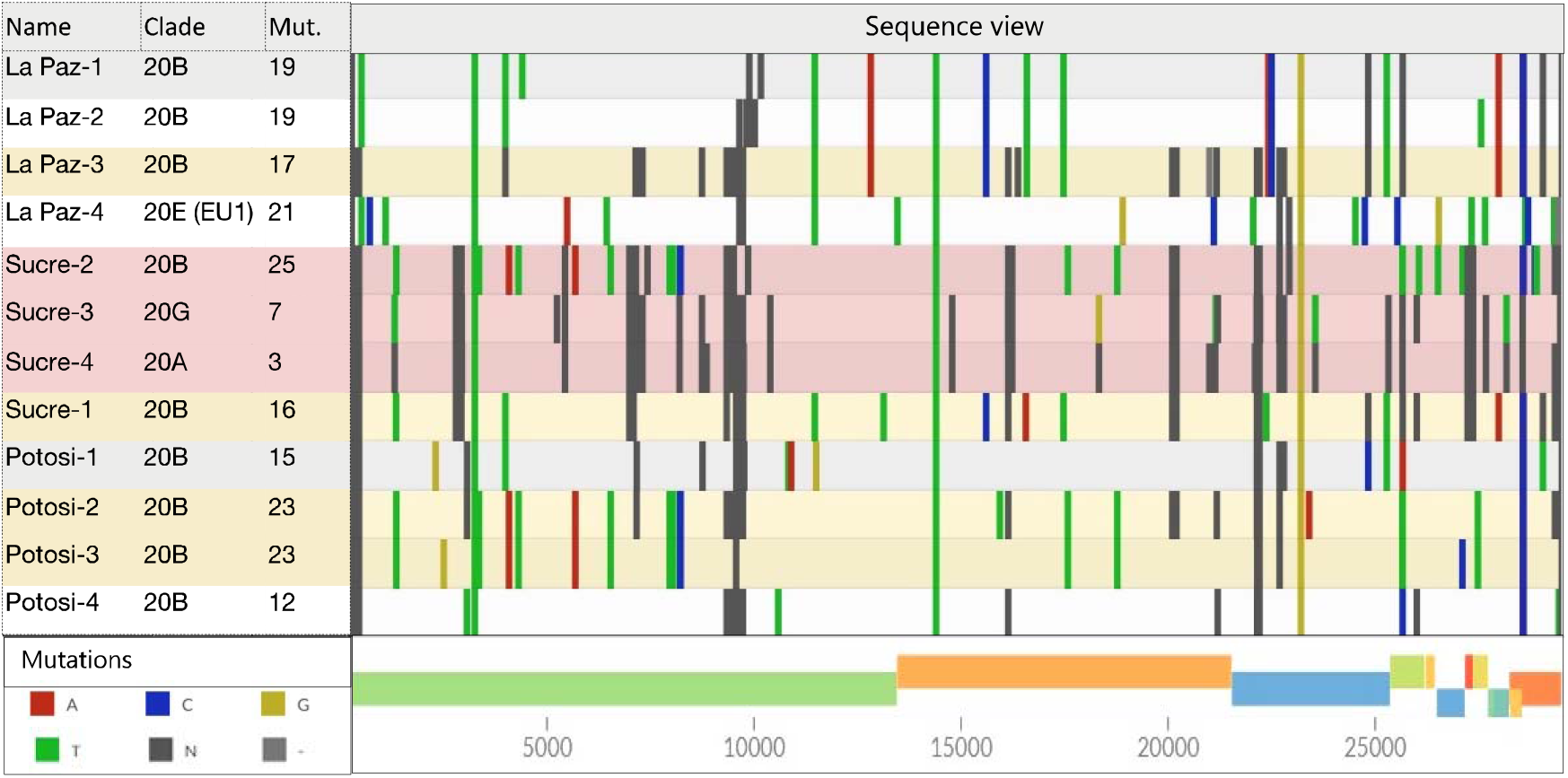
SARS-CoV-2 genome sequence visualization of samples from Bolivia. Colored line markers represent mutations from the Wuhan-Hu reference genome. Mutations color legend can be found at the left-bottom corner. Dark grey regions represent missing data.

We evaluated two annealing temperatures (65°C and 63°C) for the Multiplex PCR step given we used a different enzyme from the original protocol developed by the ARTIC network. However, the Multiplex PCR at 63°C did not amplified several regions resulting in a low genome coverage, with only 22 % coverage of the whole genome.

The library preparation was done with the Rapid PCR Barcoding kit which introduces barcode reads through tagmentation without the need of ligation. Given that several amplicons must be barcoded to cover the whole SARS-CoV-2 genome we extended the incubation time designed in the original protocol from ONT at the tagmentation step from 1 to 5 minutes. This library preparation allowed us to cover in average 98,7 % of the whole SARS-CoV-2 genome. However, the coverage was not uniform and 4 regions were not successfully amplified (Fig. 1).

### Genome sequencing of SARS-CoV-2

Samples were sequenced for 18 h. and we obtained more than 6 million reads that passed quality control and we successfully mapped a total of 3,9 million reads with an average of 329 thousand reads per sample. The average sequencing depth was 324,4X allowing to cover in average a 98,7% of the SARS-CoV-2 Wuhan-Hu reference genome (Table 1). The number of nucleotide mutations observed ranged from 3 to 25. Missing data accounted in average a 4,53% of the reads, distributed in different regions where the coverage was not deep enough to generate a consensus sequence (Fig. 1).

#### Variant identification and Phylogenomic analysis

The analyzed genomes were assigned to 4 nextstrain clades. In the city of Sucre, clades 20A, 20B and 20G were observed. In La Paz, clades 20B and 20E were observed, while in Potosi, only variants belonging to the 20B clade were observed. Further, the twelve genomes were assigned to five different lineages B.1.1, B.1.177, B.1.1.274, B.1.1.348, B.1.2. (Table 1).

In order to evaluate regional clusters in Western-central South America, we first aligned 415 genomes from Argentina, Brazil, Chile, Peru and Paraguay together with 12 samples from Bolivia. The best substitution model for phylogenomic analysis was the General Time Reversible (GTR) and with such model we created an exploratory maximun-likelihood tree with a bootstrap of 50 iterations. Clusters that included Bolivian samples together with representative genome sequences from provinces of neighboring countries were selected to further investigate the possible regional clusters. A total of 47 genomes (Table S1) were selected for constructing a new phylogenomic tree that will have a better focus of the clusters present in Bolivia. The results suggested 2 mayor clusters of SARS-CoV-2 variants circulating in Bolivia (Fig. 2). The first cluster includes 3 samples from La Paz, one from Sucre and one from Potosi, that all belong to the B.1.1348 lineage. The second cluster includes two samples from Potosi and one from Sucre, all of which belong to the lineage B.1.1.274. One sample from Potosi and one from Sucre belongs to the lineage B.1.1. The other two samples were part of external clades that involves variants not commonly present in South America, like the 20E from La Paz and the 20G from Sucre.

**Fig. 2.**
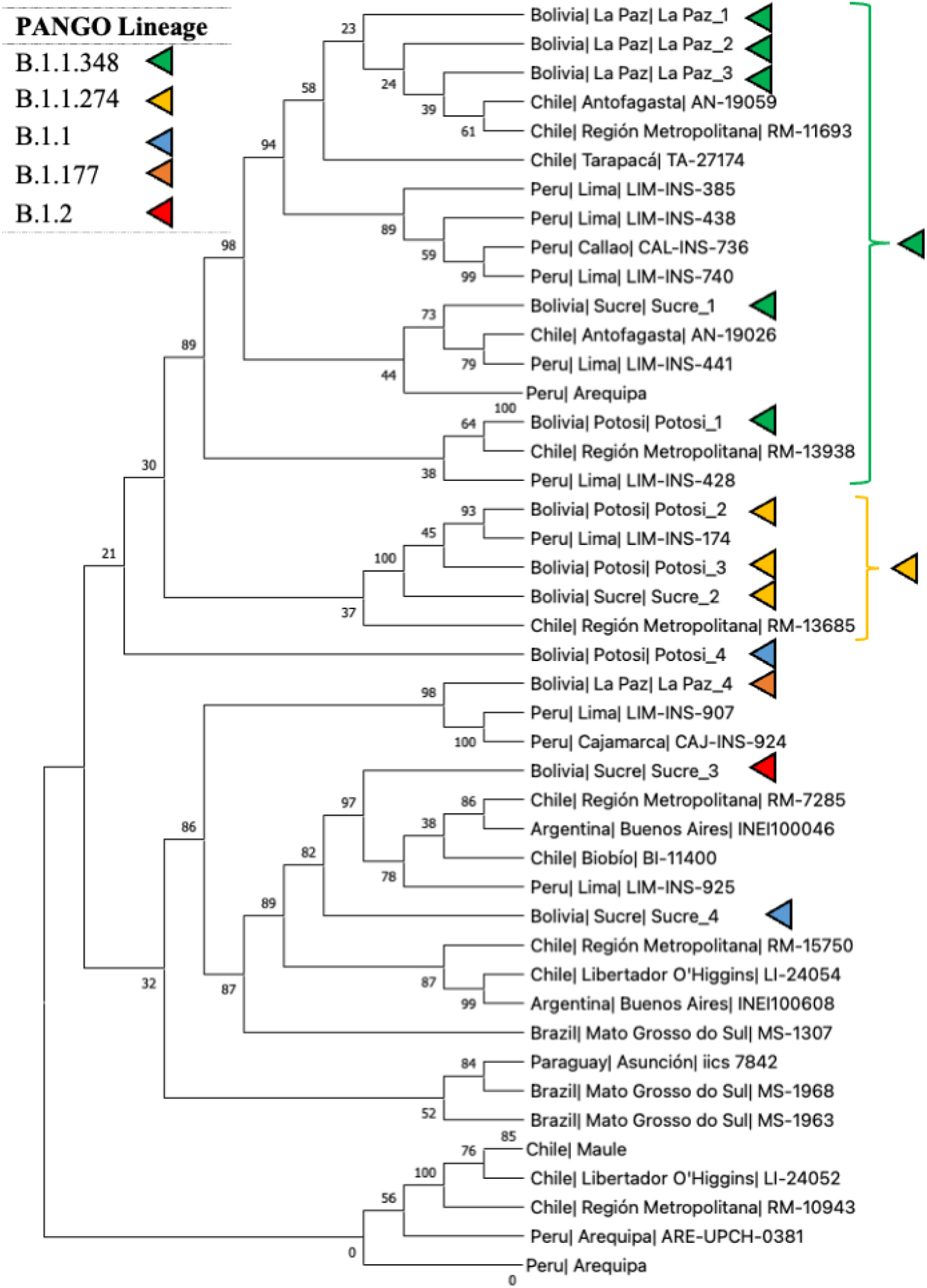
Phylogenomic relationships of SARS-CoV-2 genomes from Bolivia in January-February 2021. The evolutionary history of 12 genome sequences from Bolivia together with 35 public genomes from South America are inferred by the Maximum Likelihood method and General Time Reversible model with 1000 iterations. The two main SARS-CoV-2 clusters circulating in Bolivia are defined by curly brackets. Colors represent current PANGO lineages. Bootstrap values are given in percentage (%). Branches with two samples from the same region and country are collapsed.

## Discussions

Complete SARS-CoV-2 genomes are defined by genome lengths longer than 29.000 bp according to GISAID. Thus, we achieved to sequence complete genomes for the 12 samples analyzed. However, the number of missing data (Ns) was variable among samples. Missing data in the sequences from our workflow have two sources. The first source, which can be observed mostly in two samples from Sucre (Sucre-3 and Sucre-4) (Fig. 1), both which contain more than 3000s nucleotides missing is due to inhibition or unsuccessful amplification of certain PCR products in the Multiplex PCR step. This issue can be resolved by picking samples with higher Cts after the reverse transcription. The second source, observed in 4 regions commonly to all samples, is due to the uneven barcode insertion by Tagmentation and PCR. This uneven amplification might be due to mismatches in primer-binding sites of the barcode primers with certain DNA templates (O’Donnell et al. 2016). Another reason could be the affinity of the transposome complex with certain DNA regions because it is known that transposases have some sequence biases but the reason is not yet determined (Goryshin et al. 1998; Reznikoff 2003). This issue might be resolved by linear amplification which produces less bias and errors (Li et al. 2020) or genetically engineered transposases (Kia et al. 2017). This issue was not observed when applying library preparation with ligation instead of Tagmentation (Tyson et al. 2020).

The variant assignment results are expected because there was a predominance of the 20A and 20B (which includes the lineages B.1.1, B.1.1.274 and B.1.1.348) SARS-COV-2 variants circulating in South America (Camacho et al. 2021; Muñoz et al. 2021). The five different lineages (Table 1) found in this study are correlated with the clusters observed in the phylogenomic tree (Fig. 2). The two main clusters observed in the evolutionary analysis reveals the circulation of at least two SARS-CoV-2 lineages in Bolivia in February, 2021. Both lineages were present in Potosi and Sucre. Due to the location of Sucre, situated in between the mountainous Andean region and the lowlands we can presume that two different SARS-CoV-2 lineages were introduced independently and they evolve convergently as observed in the Metropolitan Region of Chile and Lima, the capital of Peru (Fig. 2). Although more data is needed, we can observe that only one lineage (B.1.1.348) was stablished in the governmental city La Paz. This lineage is one of the most common lineages found in South America (https://cov-lineages.org/lineages/lineage_B.1.1.348.html). Lineage B.1.1.348 accounted for 3,6% of the total SARS-CoV-2 variants of Peru until March-2021 (Camacho et al. 2021).

The other lineage (B.1.1.274) still has an unknown origin, especially in Bolivia, given that the highest prevalence reported of this lineage is in the United Kingdom (44%) and the second highest is from Russia (15%) (https://cov-lineages.org/lineages/lineage_B.1.1.274.html). The spreading of this variant reveals multiple SARS-CoV-2 introductions in Bolivia. This is expected because genomic surveillance in other south American countries have revealed this possibility before, for example in Colombia 11 different lineages were detected (Ramírez et al. 2021).

The sample from La Paz identified as 20E (EU1) is expected to be part of an external group given its presence was mostly observed in Europe (Hodcroft et al. 2021b). Similarly, sample 20G from Sucre is part of this external group because its presence was mostly observed in North America (Hodcroft et al. 2021a; Zhang et al. 2021). Both findings could be isolated events that did not spread enough to become stablished variants. However, more genomic data is needed to reach a stronger deduction.

In conclusion, the alternative library preparation for SARS-CoV-2 genome sequencing using the ARTIC protocol presented here is effective to identify SARS-CoV-2 variants with high accuracy and without the need of DNA ligation. Therefore, this can be an alternative tool to perform SARS-CoV-2 genome surveillance in developing countries.

## Additional files

Supplementary Table 1. Public genomes available from GISAID that were considered for the final dataset of the evolutionary history of SARS-CoV-2 variants circulating in Bolivia

Supplementary Table 2. Ackowledgements to all authors who deposited public genomes on GISAID that were considered for the exploratory dataset of the evolutionary history of SARS-CoV-2 circulating in Bolivia.

## Acknowledgements

We are grateful to Patricia Mollinedo, Valeria Palma and Paola Nogales of the Instituto de Investigaciones Químicas for their contributions on the development of this work. We thank BioMoLab (Former Clin&Gen Lab), La Paz, Bolivia and Laboratorio de Referencia Departamental para la Vigilancia Epidemiológica, Potosí, Bolivia. We are grateful to Josh Quick and the ARTIC Network for providing primers for the Multiplex PCR for SARS-CoV-2 sequencing and support with sequencing protocols and bioinformatic pipelines. We also thank to everyone who openly shared their genomic data on GISAID (authors listed in Supplementary Table 2).

## Contributors’ statement

OMRP and AV designed the study. OMRP, CD and AV performed the experiments. OMRP analyzed the data. OMRP and AV wrote the paper with input from all authors who reviewed and approved the final manuscript.

## Data availability

Consensus genome sequences were deposited at the Global Initiative on Sharing All Influenza Data (GISAID) repository.

