## Supplementary File 2 for "SARS-CoV-2 genome sequencing with Oxford Nanopore Technology and Rapid PCR Barcoding in Bolivia"

We gratefully acknowledge the following Authors from the Originating laboratories responsible for obtaining the specimens, as well as the Submitting laboratories where the genome data were generated and shared via GISAID, on which this research is based.

All Submitters of data may be contacted directly via www.gisaid.org

Authors are sorted alphabetically.

| Accession ID | Originating Laboratory | Submitting Laboratory | Authors |
|---|---|---|---|
| EPI_ISL_1395778, EPI_ISL_1395779, EPI_ISL_1395780 | Laboratorio de Virología del Hospital de Niños Dr. Ricardo Gutierrez | Área de Secuenciación del Laboratorio de Virología del Hospital de Niños Dr. Ricardo Gutierrez on behalf of 'Proyecto Argentino Interinstitucional de genomica de SARS-CoV-2' (PAIS Consortium) | Alexay, S; Thomas, G; Medina, C; Labarta, N; Streitenberger, C; Villegas, E; Barreda Frank, M; Grandis, E; Acevedo, ME; Alvarez Lopez, C; Jacques, O; Mistchenko, A; Nabaes Jodar, M; Goya, S; Lusso, S; Acuña, D; Natale, MI; Valinotto, LE; Viegas, M. |
| EPI_ISL_1395790 | Laboratorio de salud pública, Facultad de Ciencias Exactas, UNLP | Área de Secuenciación del Laboratorio de Virología del Hospital de Niños Dr. Ricardo Gutierrez on behalf of 'Proyecto Argentino Interinstitucional de genomica de SARS-CoV-2' (PAIS Consortium) | Rosana Isabel Toro; Andrés Angelletti; Víctoria Cabassi; Victoria Nadalich; Andrés Cordero; Carina Tersigni; Laura Delaplace; Nabaes Jodar, M; Goya, S; Lusso, S; Acuña, D; Natale, MI; Valinotto, LE; Viegas, M. |
| EPI_ISL_1395791, EPI_ISL_1395793 | Laboratorio de Virología del Hospital de Niños Dr. Ricardo Gutierrez | Área de Secuenciación del Laboratorio de Virología del Hospital de Niños Dr. Ricardo Gutierrez on behalf of 'Proyecto Argentino Interinstitucional de genomica de SARS-CoV-2' (PAIS Consortium) | Alexay, S; Thomas, G; Medina, C; Labarta, N; Streitenberger, C; Villegas, E; Barreda Frank, M; Grandis, E; Acevedo, ME; Alvarez Lopez, C; Jacques, O; Mistchenko, A; Nabaes Jodar, M; Goya, S; Lusso, S; Acuña, D; Natale, MI; Valinotto, LE; Viegas, M. |
| EPI_ISL_1395802 | Instituto de Investigaciones Biomédicas en Retrovirus y SIDA (INBIRS) | Área de Secuenciación del Laboratorio de Virología del Hospital de Niños Dr. Ricardo Gutierrez on behalf of 'Proyecto Argentino Interinstitucional de genomica de SARS-CoV-2' (PAIS Consortium) | Vanesa Seery; Federico Remes Lenicov; Horacio Salomón; Nabaes Jodar, M; Goya, S; Lusso, S; Acuña, D; Alexay, S; Natale, MI; Valinotto, LE; Viegas, M. |
| EPI_ISL_1395830, EPI_ISL_1395831, EPI_ISL_1395832 | Inmunología del Hospital Perrando e Instituto de Medicina Regional de la UNNE | Grupo de Genómica y Bioinformática del Instituto de Investigación de la Cadena Láctea CONICET-INTA on behalf of 'Proyecto Argentino Interinstitucional de genomica de SARS-CoV-2' (PAIS Consortium) | María Delia Foussal, Gerardo Deluca, Natalia Andrea Ayala, María Verónica Gómez, Gustavo Giusiano, Horacio Lucero, Marcelo Marin, Antonieta Cayré, Laura Lescano, Eberhardt, MF, Irazoqui, Amadio, AF |
| EPI_ISL_1395936, EPI_ISL_1395939 | Laboratorio Central de Salud Pública | Grupo de Genómica y Bioinformática del Instituto de Investigación de la Cadena Láctea CONICET-INTA on behalf of 'Proyecto Argentino Interinstitucional de genomica de SARS-CoV-2' (PAIS Consortium) | Natalia Andrea Ayala y María Verónica Gómez, Erica Struss, Esteban Paredes, Antonieta Cayré, Laura Lescano, Eberhardt, MF, Irazoqui, Amadio, AF |
| EPI_ISL_1395990, EPI_ISL_1395991, EPI_ISL_1395992, EPI_ISL_1396030, EPI_ISL_1396047 | Laboratorio Central, Ministerio de Salud Cordoba | Instituto de Patologia Vegetal (CIAP-INTA) on behalf of 'Proyecto Argentino Interinstitucional de genomica de SARS-CoV-2' (PAIS Consortium) | Fernández, FD; Marquez, N.; Debat, HJ.; Re, V.; Pisano, M.B.; Castro, G.; Barbas, G. |
| EPI_ISL_1396224, EPI_ISL_1396225 | Laboratorio de Virología del Hospital de Niños Dr. Ricardo Gutierrez | Área de Secuenciación del Laboratorio de Virología del Hospital de Niños Dr. Ricardo Gutierrez on behalf of 'Proyecto Argentino Interinstitucional de genomica de SARS-CoV-2' (PAIS Consortium) | Alexay, S; Thomas, G; Medina, C; Labarta, N; Streitenberger, C; Villegas, E; Barreda Frank, M; Grandis, E; Acevedo, ME; Alvarez Lopez, C; Jacques, O; Mistchenko, A; Nabaes Jodar, M; Goya, S; Lusso, S; Acuña, D; Natale, MI; Valinotto, LE; Viegas, M. |
| EPI_ISL_1396356, EPI_ISL_1396360, EPI_ISL_1396361 | Laboratorio del Hospital Regional Ushuaia Gdor. Ernesto Campos | Nodo de Secuenciación Tierra del Fuego - Hospital Regional Ushuaia - Centro Austral De Investigaciones Científicas - Universidad Nacional De Tierra Del Fuego on behalf of 'Proyecto Argentino Interinstitucional de genomica de SARS-CoV-2' (PAIS Consortium) | Carina Andrea De Roccis, Gabriel Alejandro Castro, Silvana Beatriz Cáceres, Carolina Beatriz Yulan, Manuel Fabian Boutureira, Alejandro Ezequiel Rojas, Fernando Gallego, Santiago Guillermo Ceballos, Cristina Fernanda Nardi, Ivan Dario Gramundi |
| EPI_ISL_2007525, EPI_ISL_2007526, EPI_ISL_2007545 | Laboratorio de Virología del Hospital de Niños Dr. Ricardo Gutierrez | Área de Secuenciación del Laboratorio de Virología del Hospital de Niños Dr. Ricardo Gutierrez on behalf of 'Proyecto Argentino Interinstitucional de genomica de SARS-CoV-2' (PAIS Consortium) | Alexay, S; Thomas, G; Medina, C; Labarta, N; Streitenberger, C; Villegas, E; Barreda Frank, M; Grandis, E; Acevedo, ME; Alvarez Lopez, C; Jacques, O; Mistchenko, A; Nabaes Jodar, M; Goya, S; Lusso, S; Acuña, D; Natale, MI; Valinotto, LE; Viegas, M. |
| EPI_ISL_2104813, EPI_ISL_2104814, EPI_ISL_2104815, EPI_ISL_2104819, EPI_ISL_2104824, EPI_ISL_2104825, EPI_ISL_2104826, EPI_ISL_2105566, EPI_ISL_2105567, EPI_ISL_2105568, EPI_ISL_2105569, EPI_ISL_2105570, EPI_ISL_2105571, EPI_ISL_2105572, EPI_ISL_2105573, EPI_ISL_2105574, EPI_ISL_2105575, EPI_ISL_2135135 | | | |
| see above | Servicio Virosis Respiratorias-Departamento Virología-INEI | Instituto Nacional Enfermedades Infecciosas C.G.Malbran | Baumeister E., Avaro M., Benedetti E., Russo M., Dattero ME, Pontoriero A., Cisterna D., Molina V., Perandones C., Tuduri E., Lorenzo F., Poklepovich T., Campos J. |

We gratefully acknowledge the following Authors from the Originating laboratories responsible for obtaining the specimens, as well as the Submitting laboratories where the genome data were generated and shared via GISAID, on which this research is based.

All Submitters of data may be contacted directly via www.gisaid.org

Authors are sorted alphabetically.

| Accession ID | Originating Laboratory | Submitting Laboratory | Authors |
|---|---|---|---|
| EPI_ISL_1492645, EPI_ISL_1492646, EPI_ISL_1492647, EPI_ISL_1492648, EPI_ISL_1492649, EPI_ISL_1492650, EPI_ISL_1492651, EPI_ISL_1492652 | Laboratorio Central de Salud Publica de Paraguay | Laboratorio Central de Salud Publica de Paraguay | Marta Giovanetti, María José Ortega, Andrea Gómez de la Fuente, Shirley Villalba, Juan Torales, María Liz Gamarra, Vagner Fonseca, Flavia Aburjaile, Talita Adelino, Luiz Carlos Junior Alcantara, Cynthia Vázquez |
| EPI_ISL_2234880, EPI_ISL_2234881, EPI_ISL_2234883, EPI_ISL_2234884, EPI_ISL_2234887, EPI_ISL_2234889, EPI_ISL_2234890, EPI_ISL_2234892, EPI_ISL_2234894, EPI_ISL_2234896, EPI_ISL_2234897, EPI_ISL_2234899 | | | |
| see above | IICS-UNA | IICS-UNA | Magaly Martinez, Adriana Valenzuela, Alejandra Rojas, Chyntia Diaz, Eva Nara, Fatima Cardozo, Florencia del Puerto, Joel Ortiz, Jonas Fernandez, Laura Franco, Laura Mendoza, Leticia Rojas, Maria Eugenia Galeano. |

We gratefully acknowledge the following Authors from the Originating laboratories responsible for obtaining the specimens, as well as the Submitting laboratories where the genome data were generated and shared via GISAID, on which this research is based.

All Submitters of data may be contacted directly via www.gisaid.org

Authors are sorted alphabetically.

| Accession ID | Originating Laboratory | Submitting Laboratory | Authors |
|---|---|---|---|
| EPI_ISL_1111069, EPI_ISL_1111127, EPI_ISL_1111128, EPI_ISL_1111129, EPI_ISL_1111130, EPI_ISL_1111131, EPI_ISL_1111156, EPI_ISL_1111157, EPI_ISL_1111159, EPI_ISL_1111160, EPI_ISL_1111161, EPI_ISL_1111281, EPI_ISL_1111282, EPI_ISL_1111283, EPI_ISL_1111284, EPI_ISL_1111285, EPI_ISL_1111286, EPI_ISL_1111287, EPI_ISL_1111288, EPI_ISL_1111289, EPI_ISL_1111290, EPI_ISL_1111291, EPI_ISL_1111292, EPI_ISL_1111293, EPI_ISL_1111294, EPI_ISL_1111295, EPI_ISL_1111296, EPI_ISL_1111297, EPI_ISL_1111298, EPI_ISL_1111299, EPI_ISL_1111300, EPI_ISL_1111301, EPI_ISL_1111302, EPI_ISL_1111303, EPI_ISL_1111304, EPI_ISL_1111305, EPI_ISL_1111306, EPI_ISL_1111307, EPI_ISL_1111316, EPI_ISL_1111317, EPI_ISL_1111318, EPI_ISL_1111319, EPI_ISL_1111320, EPI_ISL_1111321, EPI_ISL_1111322, EPI_ISL_1111323, EPI_ISL_1111324, EPI_ISL_1111325, EPI_ISL_1111326, EPI_ISL_1111327, EPI_ISL_1111328, EPI_ISL_1111329, EPI_ISL_1111330, EPI_ISL_1111331, EPI_ISL_1111332, EPI_ISL_1111334, EPI_ISL_1111335, EPI_ISL_1111336, EPI_ISL_1111337, EPI_ISL_1111338, EPI_ISL_1111339, EPI_ISL_1111340, EPI_ISL_1111341, EPI_ISL_1111343, EPI_ISL_1111344, EPI_ISL_1111345, EPI_ISL_1111346, EPI_ISL_1111347, EPI_ISL_1111348, EPI_ISL_1111349, EPI_ISL_1111350, EPI_ISL_1111351, EPI_ISL_1111353, EPI_ISL_1111354, EPI_ISL_1111355, EPI_ISL_1111446, EPI_ISL_1111447, EPI_ISL_1111448, EPI_ISL_1111449, EPI_ISL_1111450, EPI_ISL_1111451, EPI_ISL_1111452, EPI_ISL_1111453, EPI_ISL_1111454, EPI_ISL_1111455, EPI_ISL_1111456, EPI_ISL_1111457, EPI_ISL_1111458, EPI_ISL_1111459, EPI_ISL_1111460, EPI_ISL_1111461, EPI_ISL_1111465, EPI_ISL_1111466, EPI_ISL_1111468, EPI_ISL_1111469, EPI_ISL_1111471, EPI_ISL_1111472, EPI_ISL_1111473, EPI_ISL_1111474, EPI_ISL_1111475, EPI_ISL_1111479, EPI_ISL_1111487, EPI_ISL_1111490, EPI_ISL_1111496, EPI_ISL_1111498, EPI_ISL_1111500, EPI_ISL_1111501, EPI_ISL_1111503, EPI_ISL_1111504, EPI_ISL_1111505 | | | |
| see above | Laboratorio de Referencia Nacional de Virus Respiratorio. Instituto Nacional de Salud Perú | Laboratorio de Referencia Nacional de Enteropatógenos. Instituto Nacional de Salud del Perú | Ronnie Gavilan Chavez, Junior Caro Castro, Willi Quino Sifuentes, Veronica Hurtado Vela, Iris Silva Molina, Fiorella Orellana Peralta |
| EPI_ISL_1138413, EPI_ISL_1138414, EPI_ISL_1138415, EPI_ISL_1138416, EPI_ISL_1138417, EPI_ISL_1138418, EPI_ISL_1138419, EPI_ISL_1138420, EPI_ISL_1138421, EPI_ISL_1534670, EPI_ISL_1534671, EPI_ISL_1534673, EPI_ISL_1534674, EPI_ISL_1534675, EPI_ISL_1534686, EPI_ISL_1534687, EPI_ISL_1534688, EPI_ISL_1534689 | | | |
| see above | Laboratorio de Referencia Nacional de Virus Respiratorio. Instituto Nacional de Salud Perú | Laboratorio de Referencia Nacional de Biotecnología y Biología Molecular. Instituto Nacional de Salud Perú | Carlos Padilla Rojas, Karolyn Vega Chozo, Luis Barcena, Priscila Lope Pari, Omar Caceres Rey, Marco Galarza Perez, Maribel Huaringa Nuñez, Johanna Balbuena Torrez, Henri Bailon Calderon, Nancy Rojas Serrano |
| EPI_ISL_1548074 | Laboratorio de Referencia Nacional de Virus Respiratorio. Instituto Nacional de Salud Peru | Laboratorio de Referencia Nacional de Biotecnología y Biología Molecular. Instituto Nacional de Salud Perú | Carlos Padilla Rojas, Karolyn Vega Chozo, Luis Barcena, Priscila Lope Pari, Omar Caceres Rey, Marco Galarza Perez, Maribel Huaringa Nuñez, Johanna Balbuena Torrez, Henri Bailon Calderon, Nancy Rojas Serrano |
| EPI_ISL_1591087, EPI_ISL_1591088, EPI_ISL_1591089, EPI_ISL_1591090, EPI_ISL_1591091, EPI_ISL_1591092, EPI_ISL_1591093, EPI_ISL_1593725, EPI_ISL_1593726 | NAMRU-6 | Pathogen Discovery, Respiratory Viruses Branch, Division of Viral Diseases, Centers for Disease Control and Prevention | Yan Li, Ying Tao, Anna Kelleher, Jing Zhang, Anna Montmayeur, Brian Lynch, Krista Queen, Anna Uehara, Peter Cook, Han Jia Justin Ng, Rachel Marine, Clinton R. Paden, Haibin Wang, Mark Burroughs, Justin Lee, Adam Retchless, Suxiang Tong |
| EPI_ISL_1629762, EPI_ISL_1629763 | Hospital Carlos Alberto Seguín Escobedo - EsSalud | Laboratorio de Genómica Microbiana, Universidad Peruana Cayetano Heredia | Lenin Maturrano, Pablo Tsukayama, Alejandra Dávila-Barclay, Guillermo Salvatierra, Luis González, Pedro E. Romero, Diego Cuicapuza, Janet Huancachoque, Pool Marcos |
| EPI_ISL_1629764 | Instituto de Medicina Tropical Alexander Von Humboldt, Universidad Peruana Cayetano Heredia | Laboratorio de Genómica Microbiana, Universidad Peruana Cayetano Heredia | Lenin Maturrano, Pablo Tsukayama, Alejandra Dávila-Barclay, Guillermo Salvatierra, Luis González, Pedro E. Romero, Diego Cuicapuza, Janet Huancachoque, Pool Marcos |
| EPI_ISL_1629765, EPI_ISL_1629766, EPI_ISL_1629767, EPI_ISL_1629768, EPI_ISL_1629769, EPI_ISL_1629770, EPI_ISL_1629771, EPI_ISL_1629772, EPI_ISL_1629773, EPI_ISL_1629774, EPI_ISL_1629775, EPI_ISL_1629776, EPI_ISL_1629777, EPI_ISL_1629778, EPI_ISL_1629779, EPI_ISL_1629780, EPI_ISL_1629781, EPI_ISL_1629782 | | | |
| see above | Hospital Carlos Alberto Seguín Escobedo - EsSalud | Laboratorio de Genómica Microbiana, Universidad Peruana Cayetano Heredia | Lenin Maturrano, Pablo Tsukayama, Alejandra Dávila-Barclay, Guillermo Salvatierra, Luis González, Pedro E. Romero, Janet Huancachoque, Pool Marcos, Fernando Sánchez Fragoso, Lin Zevallos Cuarite |
| EPI_ISL_1629788, EPI_ISL_1629789 | Laboratorio de Referencia Nacional de Virus Respiratorios, Instituto Nacional de Salud Peru | Laboratorio de Genómica Microbiana, Universidad Peruana Cayetano Heredia | Lenin Maturrano, Pablo Tsukayama, Alejandra Dávila-Barclay, Guillermo Salvatierra, Luis González, Pedro E. Romero, Diego Cuicapuza, Janet Huancachoque, Pool Marcos, Maribel Huaringa, Priscilla Lope, Nancy Rojas |
| EPI_ISL_1629790 | Laboratorio de Referencia Nacional de Virus Respiratorios, Instituto Nacional de Salud Peru | Laboratorio de Genómica Microbiana, Universidad Peruana Cayetano Heredia | Lenin Maturrano, Pablo Tsukayama, Alejandra Dávila-Barclay, Guillermo Salvatierra, Luis González, Pedro E. Romero, Diego Cuicapuza, Janet Huancachoque, Pool Marcos |
| EPI_ISL_1701773, EPI_ISL_1701774 | Laboratorio de Referencia Nacional de Virus Respiratorio. Instituto Nacional de Salud Perú | Laboratorio de Referencia Nacional de Biotecnología y Biología Molecular. Instituto Nacional de Salud Perú | Carlos Padilla Rojas, Karolyn Vega Chozo, Luis Barcena, Priscila Lope Pari, Omar Caceres Rey, Marco Galarza Perez, Maribel Huaringa Nuñez, Johanna Balbuena Torrez, Henri Bailon Calderon, Nancy Rojas Serrano |
| EPI_ISL_925173 | Laboratorio de Referencia Nacional de Enteropatógenos. Instituto Nacional de Salud del Perú | Laboratorio de Referencia Nacional de Enteropatógenos. Instituto Nacional de Salud del Perú | Ronnie Gavilan Chavez, Junior Caro Castro, Willi Quino Sifuentes, Veronica Hurtado Vela, Iris Silva Molina, Fiorella Orellana Peralta, |
| EPI_ISL_943570 | Laboratorio de Referencia Nacional de Virus Respiratorio. Instituto Nacional de Salud Perú | Laboratorio de Referencia Nacional de Biotecnología y Biología Molecular. Instituto Nacional de Salud Perú | Carlos Padilla Rojas, Karolyn Vega Chozo, Luis Barcena, Priscila Lope Pari, Omar Caceres Rey, Marco Galarza Perez, Maribel Huaringa Nuñez, Johanna Balbuena Torrez, Henri Bailon Calderon, Nancy Rojas Serrano |

We gratefully acknowledge the following Authors from the Originating laboratories responsible for obtaining the specimens, as well as the Submitting laboratories where the genome data were generated and shared via GISAID, on which this research is based.

All Submitters of data may be contacted directly via www.gisaid.org

Authors are sorted alphabetically.

| Accession ID | Originating Laboratory | Submitting Laboratory | Authors |
|---|---|---|---|
| EPI_ISL_1167668, EPI_ISL_1167669, EPI_ISL_1167670, EPI_ISL_1167674, EPI_ISL_1167675, EPI_ISL_1167676, EPI_ISL_1167694, EPI_ISL_1167695, EPI_ISL_1167735, EPI_ISL_1167759, EPI_ISL_1167760, EPI_ISL_1167761, EPI_ISL_1167762, EPI_ISL_1167763, EPI_ISL_1167764, EPI_ISL_1167765, EPI_ISL_1167767, EPI_ISL_1167768, EPI_ISL_1167769, EPI_ISL_1167771, EPI_ISL_1167772, EPI_ISL_1167775, EPI_ISL_1167776, EPI_ISL_1167777, EPI_ISL_1167778, EPI_ISL_1167804, EPI_ISL_1167839, EPI_ISL_1167857, EPI_ISL_1167858, EPI_ISL_1167865, EPI_ISL_1167867, EPI_ISL_1167871, EPI_ISL_1167872, EPI_ISL_1167874, EPI_ISL_1167875, EPI_ISL_1167876, EPI_ISL_1167884, EPI_ISL_1167886, EPI_ISL_1167888, EPI_ISL_1167890, EPI_ISL_1167891, EPI_ISL_1167893, EPI_ISL_1167896, EPI_ISL_1167897, EPI_ISL_1167899, EPI_ISL_1167901, EPI_ISL_1167902, EPI_ISL_1167903, EPI_ISL_1167906, EPI_ISL_1167909, EPI_ISL_1167912, EPI_ISL_1167915, EPI_ISL_1167918, EPI_ISL_1167922, EPI_ISL_1167927, EPI_ISL_1167929, EPI_ISL_1167931, EPI_ISL_1300479, EPI_ISL_1300480, EPI_ISL_1300481, EPI_ISL_1300500, EPI_ISL_1300501, EPI_ISL_1300511, EPI_ISL_1321453, EPI_ISL_1321454, EPI_ISL_1321468, EPI_ISL_1321476, EPI_ISL_1321492, EPI_ISL_1321495, EPI_ISL_1321498, EPI_ISL_1321506, EPI_ISL_1321509, EPI_ISL_1321528, EPI_ISL_1321531, EPI_ISL_1321532, EPI_ISL_1321533, EPI_ISL_1321534, EPI_ISL_1321535, EPI_ISL_1321536, EPI_ISL_1321563, EPI_ISL_1321569, EPI_ISL_1321570, EPI_ISL_1321571, EPI_ISL_1321572, EPI_ISL_1321573, EPI_ISL_1321574, EPI_ISL_1321575, EPI_ISL_1321576, EPI_ISL_1321577, EPI_ISL_1321578, EPI_ISL_1321579, EPI_ISL_1321580, EPI_ISL_1321581, EPI_ISL_1321582, EPI_ISL_1321583, EPI_ISL_1321584, EPI_ISL_1470483, EPI_ISL_1470484, EPI_ISL_1470485, EPI_ISL_1470486, EPI_ISL_1470487, EPI_ISL_1470488, EPI_ISL_1470489 | | | |
| see above | Genetica Molecular and Subdepartamento de Virologia ISP Chile | Instituto de Salud Publica de Chile | Javier Tognarelli, Karen Orostica, Barbara Parra, Loredana Arata, Jaime Lagos, Gisselle Barra, Patricia Bustos, Rodrigo Fasce, Andres Castillo, Jorge Fernandez |
| EPI_ISL_2009615 | Genetica Molecular and Subdepartamento de Virologia ISP Chile | Instituto de Salud Publica de Chile | Javier Tognarelli, Karen Orostica, Barbara Parra, Loredana Arata, Gisselle Barra, Patricia Bustos, Rodrigo Fasce, Andres Castillo, Soledad Ulloa, Jorge Fernandez |
| EPI_ISL_2391363, EPI_ISL_2391364, EPI_ISL_2391365 | Genetica Molecular and Subdepartamento de Virologia ISP Chile | Instituto de Salud Publica de Chile | Karen Orostica, Constanza Campano, Barbara Parra, Loredana Arata, Gisselle Barra, Patricia Bustos, Rodrigo Fasce, Javier Tognarelli, Andres Castillo, Soledad Ulloa, Jorge Fernandez |
| EPI_ISL_2728580 | Laboratorio de Infectologia y Virologia Molecular | Laboratory of Molecular Virology, School of Medicine, Pontificia Universidad Catolica de Chile | Catalina Pardo-Roa, Constanza Martinez-Valdevenito, Jennifer Angulo, Tamara Garcia-Salum, Estefany Poblete, Maria Jose Avendano, Leonardo I. Almonacid, Ana Maria Contreras, Carlos Palma, Jorge Levican, Eileen Serrano, Constanza Maldonado, M. Belen Leyton, Erick Salinas, Andres E. Munoz-Marcos, Francisco Melo, Marcela Ferres, Rafael A. Medina |

We gratefully acknowledge the following Authors from the Originating laboratories responsible for obtaining the specimens, as well as the Submitting laboratories where the genome data were generated and shared via GISAID, on which this research is based.

All Submitters of data may be contacted directly via www.gisaid.org

Authors are sorted alphabetically.

| Accession ID | Originating Laboratory | Submitting Laboratory | Authors |
|---|---|---|---|
| EPI_ISL_2245062, EPI_ISL_2248781 | Laboratório Central de Saúde Pública do Acre | Coordenação Geral de Laboratórios de Saúde Pública (CGLAB/DAEVS/SVS/MS) | Vagner Fonseca, et al. |
| EPI_ISL_2612315, EPI_ISL_2612317, EPI_ISL_2612321, EPI_ISL_2612322, EPI_ISL_2612323, EPI_ISL_2612324, EPI_ISL_2612336, EPI_ISL_2612337, EPI_ISL_2612345, EPI_ISL_2612346, EPI_ISL_2612347, EPI_ISL_2612348, EPI_ISL_2612349, EPI_ISL_2612350, EPI_ISL_2612351, EPI_ISL_2612352, EPI_ISL_2612353, EPI_ISL_2612354, EPI_ISL_2612355, EPI_ISL_2612356, EPI_ISL_2612357, EPI_ISL_2612358, EPI_ISL_2612359, EPI_ISL_2612360, EPI_ISL_2612361, EPI_ISL_2612362, EPI_ISL_2612363, EPI_ISL_2612364, EPI_ISL_2612393, EPI_ISL_2612394, EPI_ISL_2612395, EPI_ISL_2612396, EPI_ISL_2612404, EPI_ISL_2612407 | | | |
| see above | Centro de Infectologia Charles Mérieux/ Laboratório Rodolphe Mérieux, FUNDHACRE | Bioinformatics Laboratory / LNCC | Alessandra P Lamarca, Luiz G P de Almeida, Ronaldo da Silva F Jr,Douglas Terra Machado, Alexandra L Gerber, Ana Paula de C Guimarães, Cirley Maria de Oliveira Lobato, Andreas Stocker, Luiz Fellype Alves de Souza, Ana Tereza R Vasconcelos |

We gratefully acknowledge the following Authors from the Originating laboratories responsible for obtaining the specimens, as well as the Submitting laboratories where the genome data were generated and shared via GISAID, on which this research is based.

All Submitters of data may be contacted directly via www.gisaid.org

Authors are sorted alphabetically.

| Accession ID | Originating Laboratory | Submitting Laboratory | Authors |
|---|---|---|---|
| EPI_ISL_1293052, EPI_ISL_1293053, EPI_ISL_1293054, EPI_ISL_1293055, EPI_ISL_1303499, EPI_ISL_1303500, EPI_ISL_1303501, EPI_ISL_1303502, EPI_ISL_1303503, EPI_ISL_1303504, EPI_ISL_1303505 | | | |
| see above | LACEN de Rondonia | Instituto Adolfo Lutz, Interdiciplinary Procedures Center, Strategic Laboratory | Claudio Tavares Sacchi, Claudia Regina Gonçalves, Erica Valessa Ramos Gomes, Karoline Rodrigues Campos, Caio Vinicius Dias Lopes |
| EPI_ISL_1493583, EPI_ISL_1493595, EPI_ISL_1493596, EPI_ISL_1493597, EPI_ISL_1493598, EPI_ISL_1493599, EPI_ISL_1493600, EPI_ISL_1520107 | LACEN do Estado de Rondonia | Instituto Adolfo Lutz, Interdiciplinary Procedures Center, Strategic Laboratory | Claudio Tavares Sacchi, Claudia Regina Gonçalves, Erica Valessa Ramos Gomes, Karoline Rodrigues Campos, Caio Vinicius Dias Lopes |

We gratefully acknowledge the following Authors from the Originating laboratories responsible for obtaining the specimens, as well as the Submitting laboratories where the genome data were generated and shared via GISAID, on which this research is based.

All Submitters of data may be contacted directly via www.gisaid.org

Authors are sorted alphabetically.

| Accession ID | Originating Laboratory | Submitting Laboratory | Authors |
| --- | --- | --- | --- |
| EPI_ISL_1040824 | LACEN do Mato Grosso do Sul | Instituto Adolfo Lutz, Interdiciplinary Procedures Center, Strategic Laboratory | Claudio Tavares Sacchi, Claudia Regina Gonçalves, Erica Valessa Ramos Gomes, Karoline Rodrigues Campos |
| EPI_ISL_1133274, EPI_ISL_1133279 | Laboratório Hermes Pardini | Laboratório de Biologia Integrativa, Instituto de Ciências Biológicas, Universidade Federal de Minas Gerais | Filipe Romero Rebello Moreira, Diego Menezes Bonfim, Danielle Alves Gomes Zauli, Joice do Prado Silva, Aline Brito de Lima, Frederico Scott Varella Malta, Alessandro Clayton de Souza Ferreira, Victor Cavalcanti Pardini, Daniel Costa Queiroz, Rafael Marques de Souza, Victor Emmanuel Viana Geddes, Walyson Coelho Costa, Wagner Carlos Santos Magalhaes, Rennan Garcias Moreira, Carolina Moreira Voloch, Renan Pedra de Souza, Renato Santana Aguiar. |
| EPI_ISL_1139059, EPI_ISL_1196282, EPI_ISL_1358310, EPI_ISL_1358311, EPI_ISL_1358312, EPI_ISL_1358313, EPI_ISL_1358314, EPI_ISL_1358315, EPI_ISL_1358316, EPI_ISL_1358317, EPI_ISL_1381066, EPI_ISL_1381067, EPI_ISL_1468436, EPI_ISL_1468437, EPI_ISL_1468438, EPI_ISL_1468439, EPI_ISL_1468440, EPI_ISL_1468441 | | | |
| see above | LACEN do Mato Grosso do Sul | Instituto Adolfo Lutz, Interdiciplinary Procedures Center, Strategic Laboratory | Claudio Tavares Sacchi, Claudia Regina Gonçalves, Erica Valessa Ramos Gomes, Karoline Rodrigues Campos, Caio Vinicius Dias Lopes |
| EPI_ISL_2245194 | Laboratório Central de Saúde Pública do Mato Grosso do Sul | Coordenação Geral de Laboratórios de Saúde Pública (CGLAB/DAEVS/SVS/MS) | Vagner Fonseca, et al. |
| EPI_ISL_2344240, EPI_ISL_2344241 | Instituto Butantan | Instituto de Medicina Tropical de Sao Paulo | Brazil-UK Centre for Arbovirus Discovery Diagnosis Genomics and Epidemiology (CADDE) Genomic Network - Instituto de Medicina Tropical |

We gratefully acknowledge the following Authors from the Originating laboratories responsible for obtaining the specimens, as well as the Submitting laboratories where the genome data were generated and shared via GISAID, on which this research is based.

All Submitters of data may be contacted directly via www.gisaid.org

Authors are sorted alphabetically.

| Accession ID | Originating Laboratory | Submitting Laboratory | Authors |
|---|---|---|---|
| EPI_ISL_1040824 | LACEN do Mato Grosso do Sul | Instituto Adolfo Lutz, Interdiciplinary Procedures Center, Strategic Laboratory | Claudio Tavares Sacchi, Claudia Regina Gonçalves, Erica Valessa Ramos Gomes, Karoline Rodrigues Campos |
| EPI_ISL_1139059, EPI_ISL_1196282, EPI_ISL_1358310, EPI_ISL_1358311, EPI_ISL_1358312, EPI_ISL_1358313, EPI_ISL_1358314, EPI_ISL_1358315, EPI_ISL_1358316, EPI_ISL_1358317, EPI_ISL_1381066, EPI_ISL_1381067, EPI_ISL_1468436, EPI_ISL_1468437, EPI_ISL_1468438, EPI_ISL_1468439, EPI_ISL_1468440, EPI_ISL_1468441 | | | |
| see above | LACEN do Mato Grosso do Sul | Instituto Adolfo Lutz, Interdiciplinary Procedures Center, Strategic Laboratory | Claudio Tavares Sacchi, Claudia Regina Gonçalves, Erica Valessa Ramos Gomes, Karoline Rodrigues Campos, Caio Vinicius Dias Lopes |
| EPI_ISL_2245194 | Laboratório Central de Saúde Pública do Mato Grosso do Sul | Coordenação Geral de Laboratórios de Saúde Pública (CGLAB/DAEVS/SVS/MS) | Vagner Fonseca, et al. |
